# Effects of temperature on mating behaviour and mating success: a meta-analysis

**DOI:** 10.1101/2022.05.11.491542

**Authors:** Natalie Pilakouta, Anaїs Baillet

**Author notes:** Corresponding author: Natalie Pilakouta, address: Zoology Building, University of Aberdeen, AB24 2TZ, UK).

## Abstract

1. In light of global climate change, there is a pressing need to understand how populations will respond to rising temperatures. Understanding the effects of temperature changes on mating behaviour is particularly important, given its implications for population viability.
2. To this end, we performed a meta-analysis of 53 studies to examine how temperature changes influence mating latency, choosiness, and mating success. We hypothesized that if higher temperatures make mate searching and mate assessment more costly due to an elevated metabolism, this may lead to a reduction in mating latency and choosiness, thereby increasing overall mating success.
3. We found no evidence for a global effect of temperature on mating latency, choosiness, or mating success. There was an increase in mating success when animals were exposed to higher temperatures during mating, but not when they were exposed before mating.
4. Interestingly, in a subset of studies that measured both mating latency and mating success, there was a strong negative relationship between the effect sizes for these traits. This suggests that a decrease in mating latency at higher temperatures was associated with an increase in mating success and vice versa.
5. In sum, our meta-analysis provides new insights into the effects of temperature on mating patterns. The absence of a consistent directional effect of temperature on mating behaviours and mating success suggests it may be difficult to predict changes in the strength of sexual selection in natural populations in a warming world. Nevertheless, there is some evidence that (*i*) higher temperatures during mating may lead to an increase in mating success and that (*ii*) an increase in mating success is associated with a decrease in mating latency.

## INTRODUCTION

In light of global climate change, there is a pressing need to understand how populations will respond and adapt to rising temperatures (Crozier & Hutchings 2014). Because animal behaviour is particularly labile, there is a growing body of literature investigating the effects of temperature on a wide range of behavioural traits (Abram et al. 2017). Understanding how temperature changes might affect mating behaviour and mating success is particularly important, given its link to population viability and performance (Candolin & Heuschele 2008).

A number of studies have already shown that strong sexual selection can increase population fitness and reduce the risk of extinction (Moller and Alatalo 1999, Lorch et al. 2003, Price et al. 2010, Lumley et al. 2015, Cally et al. 2019; but see Tanaka 1996). Sexual selection could therefore play a major role in the capacity of populations to cope with climate change if stronger mate preferences for ‘good genes’ can lead to higher-quality offspring (Candolin & Heuschele 2008, Martinossi-Alliberti et al. 2019, Godwin et al. 2020). In order to better understand the link between temperature and the strength and direction of sexual selection, we need to focus on reproductive traits involved in precopulatory and postcopulatory processes to identify the specific underlying mechanisms influenced by temperature variation (García-Roa et al. 2020).

Here, we performed a meta-analysis examining how an increase in temperature influences mating latency, choosiness, and mating success. We focused on plastic, rather than evolutionary, responses to changes in temperature because there are far fewer studies on the latter. Temperature can have both direct and indirect effects on mating behaviour and mating success (reviewed in Garcia-Roa et al. 2020). For example, temperature is a key determinant of metabolic rate and locomotor performance, particularly in ectotherms (Gibert et al. 2007, Lachenicht et al. 2010). In turn, metabolic rate is closely linked to activity levels (Kearney et al. 2010, Gunderson & Leal 2015) and can thus influence mate searching and spatio-temporal distributions of the two sexes (Garcia-Roa et al. 2020). As a result, temperature has been shown to modulate a wide range of precopulatory mating behaviours, including mating latency, mate choice, courtship behaviour, remating rate, and the intensity of intrasexual competition (e.g., Kvarnemo 1998, Jiao et al. 2009, Katsuki & Miyatake 2009, Conrad et al. 2017, Gudka & Santos 2019).

Two possible outcomes for our meta-analysis were as follows: (*i*) if higher temperatures make mate searching and mate assessment more energetically costly due to an elevated metabolic rate, this might lead to a reduction in mating latency and cause females to be less choosy, thereby indirectly increasing overall mating success; (*ii*) alternatively, if the benefits of mate choice are higher under warmer conditions, we might expect an increase in mating latency and choosiness, along with a decrease in mating success at the population level. In the former scenario, a reduction in mating latency and choosiness would lead to weaker sexual selection, whereas in the latter scenario an increase in mating latency and choosiness would lead to stronger sexual selection. Stronger sexual selection may in turn improve population viability by purging deleterious alleles in males that are also deleterious in females (Whitlock & Agrawal 2009, McGuigan et al. 2011).

Our meta-analysis also examined whether the relationship between temperature and mating behaviour or mating success depends on the magnitude of the temperature change, the life stage during which the temperature change occurred, and whether there was a short-term or long-term change in temperature. We expected stronger effects of temperature on mating behaviour and mating success when the change in temperature was larger and when it represented a long-term change in thermal conditions. We might also expect stronger effects when the change in temperature occurred during mating than before mating, because it would directly affect metabolism, activity levels, and locomotor performance during mate searching and assessment (Garcia-Roa et al. 2020). On the other hand, temperature changes in early development may have a more pronounced effect on mating behaviour than temperature changes in adulthood, since ‘critical windows’ in early life can affect brain development with long-term consequences for behaviour (e.g., Adewale et al. 2011, O’Connor et al. 2021).

The ability to generate general predictions for how populations will respond to climate change is crucial for management and conservation efforts, but it is difficult for individual empirical studies to address this issue, as they generally focus on a single species (Crozier & Hutchings 2014). We suggest that meta-analytical approaches present a powerful tool for better understanding and predicting the effects of rising temperatures on natural populations. By synthesizing data from published studies, meta-analyses can detect common patterns across species (Harrison 2011). Thus, in the context of predicting population responses to climate change, if a meta-analysis reveals consistent directional effects of temperature on a particular trait across a wide range of taxonomic groups, it may allow us to generate some general predictions.

## METHODS

### Search protocol and data collection

This meta-analysis was conducted following the preferred reporting items for systematic reviews (PRISMA) approach (Moher et al. 2009; Supplementary Figure 1). Database searches were conducted using Web of Science and Scopus on the 20^th^ and 21^st^ of February 2020, respectively. We used search terms that would identify studies focusing on temperature variation and mating latency, choosiness, or mating success: (“temperature” OR “thermal” OR “warm*” OR “cold*”) AND (“latency” OR “mate” or “mating” OR “mate choice” OR “choosy” OR “choosiness” OR “mat* preference” OR “copulat*” or “mating success”). All papers identified through these searches were checked for relevance based on the title and abstract (Supplementary Figure 1). After removing papers that were clearly not relevant to our question, we screened the full text of the remaining papers to find studies that contained information on mating latency, choosiness, and/or mating success under two or more temperature conditions. We considered mating latency to be a measure of individuals’ propensity to mate, which was described using different terms across papers (e.g., time to copulation, time to mating, premating period, precopulation period). We defined choosiness as “the change in mating propensity in response to alternative stimuli” (Reinhold & Schielzeth 2015). Choosiness thus referred to the strength of the mating preference when individuals were choosing between two potential mates (e.g., small vs large) or sexual signals (e.g., mating calls). Mating success referred to the proportion of experimental pairs that engaged in copulation during the mating trial.

**Figure 1.**
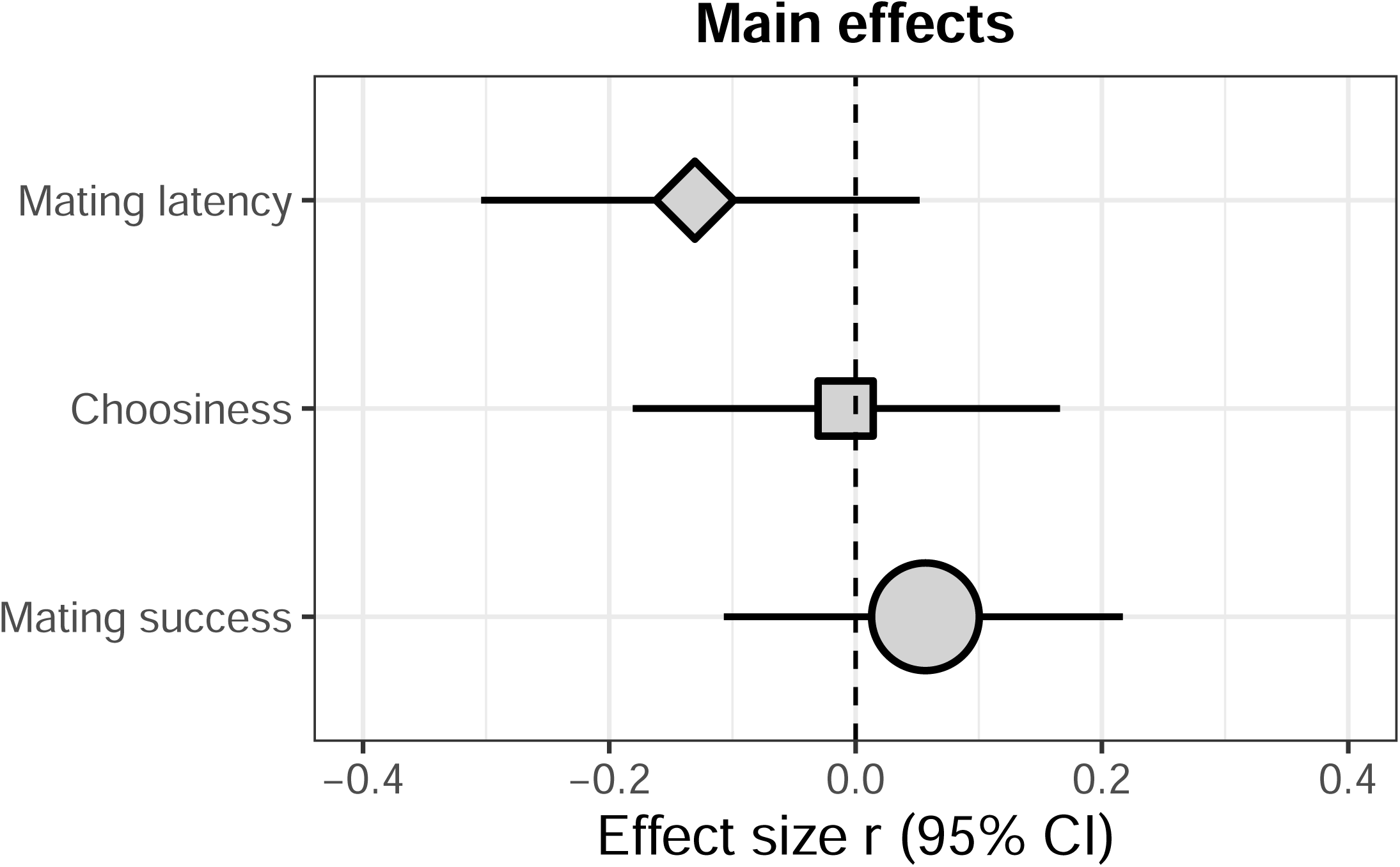
Mean effect size estimates derived from multilevel intercept-only meta-analytic models examining the effects of temperature on mating latency (diamond), choosiness (square), and mating success (circle). The relative size of each symbol represents the number of effect sizes included in that dataset (mating latency = 29, choosiness = 29, mating success = 58).

We included experimental studies with either short-term or long-term exposure to different constant temperatures within a generation. Because our aim was to examine plastic, rather than evolutionary, responses to temperature, we excluded experimental evolution studies where organisms were exposed to contrasting thermal environments over multiple generations. We also excluded studies where there were confounding variables, for example when the temperature treatments were coupled with other factors (e.g., comparisons between a warm and dry environment versus a cold and wet environment). We excluded studies where animals were exposed to fluctuating temperatures. For studies on mating latency and choosiness, we only included those where the choosing sex (usually the female) was subjected to a temperature treatment (Supplementary Table 1). For studies on mating success, we included studies where the male, female, or both were subjected to a temperature treatment (Supplementary Table 1).

**Table 1.** Number of studies (*n*), species, and effect sizes (*k*) used in our meta-analysis on the effects of temperature on mating latency, choosiness (strength of preference), and mating success. We also show the breakdown by taxonomic group for each trait.

|  | Sample sizes |  |  |
| --- | --- | --- | --- |
| | Studies ($n$) | Species | Effect sizes ($k$) |
| **Mating latency** | **19** | **14** | **29** |
| Arachnid | 1 | 1 | 1 |
| Insect | 17 | 12 | 27 |
| Mollusc | 1 | 1 | 1 |
| **Choosiness** | **14** | **14** | **29** |
| Amphibian | 1 | 1 | 1 |
| Bird | 2 | 2 | 2 |
| Crustacean | 1 | 1 | 2 |
| Fish | 2 | 2 | 5 |
| Insect | 7 | 7 | 18 |
| Reptile | 1 | 1 | 1 |
| **Mating success** | **31** | **28** | **58** |
| Collembola | 1 | 1 | 1 |
| Fish | 1 | 1 | 2 |
| Insect | 29 | 26 | 55 |

Our full-text screening also included a small number of additional references that were not identified through our literature search and were instead obtained from other sources, such as a request for relevant papers from colleagues on Twitter (*n* = 10). Data from two of these papers were deemed relevant and included in the final analysis. Overall, full-text screening identified 62 studies that met the experimental design criteria for inclusion in our meta-analysis (Supplementary Figure 1).

### Effect size calculation

To calculate effect sizes, we extracted data from the main text, tables, or figures using the image analysis software WebPlotDigitzer (Rohatgi 2019). We were unable to extract appropriate effect sizes from 13 studies due to missing test statistics or sample sizes. In these cases, we contacted the corresponding authors of these studies using a standardized email asking for the missing information. Seven of these authors responded to our email, and of those, four were able to provide the information needed to calculate effect sizes.

For each of the studies included in our meta-analysis (*n* = 53), we calculated *r* effect sizes (correlation coefficient). In our analysis, a positive *r* effect size indicates that temperature and the trait of interest are positively correlated (e.g., higher temperatures are associated with increased mating success). Some studies included multiple effect sizes for the same or different species; to control for this, we included “study” as a random factor in our analysis (see *Data analysis* for details). Overall, we collected 29 effect sizes for mating latency, 29 effect sizes for choosiness, and 58 effect sizes for mating success (Table 1). These data comprised both vertebrates (amphibians, birds, fishes, reptiles) and invertebrates (arachnids, crustaceans, insects, molluscs) with insects being the most common taxonomic group studied (Table 1).

### Data analysis

All statistical analyses were performed in R version 3.6.1 (R Core Team 2019), and figures were generated using ggplot2 (Wickham 2016). Meta-analyses were performed using Fisher’s *z* transform of the correlation coefficient (*Zr*). We then converted mean effect size estimates derived from our statistical models back to *r* for presentation in figures. Assumptions of linear mixed models (i.e., normality of residuals) were met for all models reported below.

#### Main effects models

Using the function ‘rma.mv’ from the R package ‘metafor’ (Viechtbauer 2010), we run a multilevel intercept-only meta-analytic model for each of our three traits of interest (mating latency, choosiness, and mating success) to test for an overall effect of temperature. We ran both phylogenetic and non-phylogenetic models to examine whether the evolutionary relationships between species influenced this overall effect. For the phylogenetic models, we first used the R package ‘rotl’ (Michonneau et al. 2016, OpenTree et al. 2021) to generate phylogenetic trees of the species included in our meta-analysis (Supplementary Figure 2). This tree was then imported into the ‘ape’ package (Paradis et al. 2004), and a correlation matrix obtained using the ‘vcv’ function. The resulting correlation matrix was then included in our multivariate meta-analytic models as a random factor. We included additional random effects in our phylogenetic and non-phylogenetic models to account for non-independence due to the extraction of multiple effect sizes from the same study (study ID) and included a unit level random effect (effect size ID) as a measure of residual heterogeneity (Santos et al. 2011). In non-phylogenetic models, we also included a random effect to account for the use of the same species across studies (species ID).

**Figure 2.**
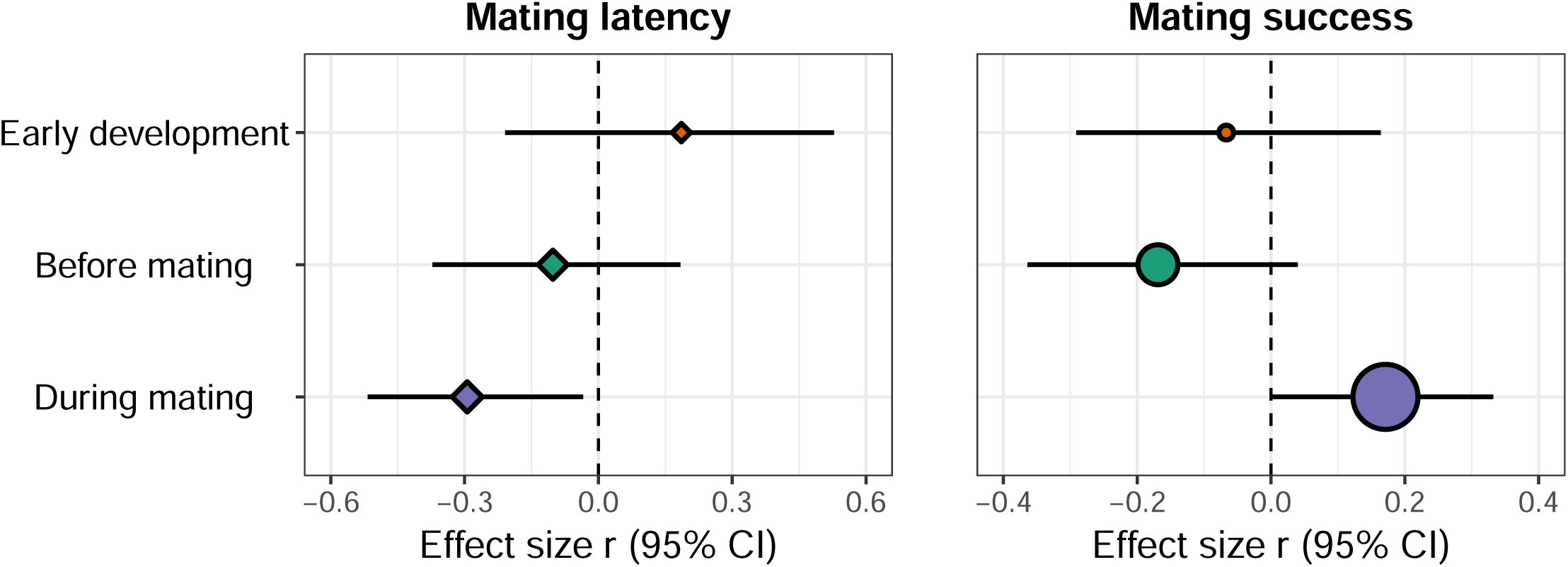
Mean effect size estimates derived from multilevel meta-regression models examining how the “time of temperature treatment” moderator (early development, adulthood before mating, or adulthood during mating) influences the relationship between temperature and mating latency (diamond) or mating success (circle). The relative size of each symbol represents the number of effect sizes included in that dataset (mating latency: early development = 6, before mating = 11, during mating = 12; mating success: early development = 6, before mating = 14, during mating = 38).

#### Moderator effects models

Because phylogeny failed to resolve any of the heterogeneity, we did not include it in our moderator effects models (Santos et al. 2011; but see Supplementary Table 2). Moderators were tested using non-phylogenetic multilevel meta-regression models with study ID, species ID, and effect size ID as random effects (Santos et al. 2011). We first included a continuous moderator (“intensity of temperature treatment”) to test whether variation in effect sizes could be explained by the extent of the temperature differences (°C) between treatments within each study. A second categorical moderator (“time of temperature treatment”) tested for differences between studies that exposed animals in early development, in adulthood before the mating trial, or during the mating trial. A third categorical moderator (“type of temperature treatment”) tested for differences between studies that exposed animals to a short-term versus a long-term temperature treatment (i.e., acute exposure versus acclimation). We defined long-term temperature treatment as any exposure to a different temperature that lasted more than 24 hr. Lastly, for mating success, we included an additional categorical moderator (“sex exposed to temperature treatment”) to test for differences between studies that exposed males, females, or both sexes to the temperature treatment. For each of these models, we calculated *R*^2^_marginal_, which describes the percentage of heterogeneity that was explained by the inclusion of a moderator (i.e., the estimated percentage decrease in heterogeneity between the main effects model and moderator model).

#### *Relationship between Zr*_mating latency_ *and Zr*_mating success_

We also examined the relationship between effect sizes (*Zr*) for mating latency and mating success from studies that measured both traits. This was the case for 14 effect sizes from 10 studies on 9 different species. We analysed this reduced dataset using (*i*) a Pearson’s correlation test between *Zr*_mating latency_ and *Zr*_mating success_, as well as (*ii*) a meta-regression with *Zr*_mating success_ as the response variable, *Zr*_mating latency_ as the moderator, and study ID, species ID, and effect size ID as random effects.

#### Publication bias tests

To examine the potential for underreporting of non-significant results, we used the function ‘regrest’ to test for funnel plot asymmetry in our meta-regression models (Nakagawa et al. 2017). We also tested for time-lag bias using (*i*) a rank correlation test between effect size and publication year for each study and (*ii*) a meta-regression with publication year as a continuous moderator (Jennions & Møller 2002).

## RESULTS

We present mean effect size estimates derived from the statistical models with 95% confidence intervals in square brackets.

### Publication bias tests

There was evidence for funnel asymmetry for mating success (*z* = -2.48, *P* = 0.013) but not for mating latency (*z* = 1.21, *P* = 0.23) or choosiness (*z* = 1.01, *P* = 0.31). This suggests a potential for publication bias in the mating success dataset (Supplementary Figure 3). In addition, our models with publication year as a continuous moderator showed a time-lag bias in the mating latency dataset (*Zr* = 0.03 [0.003, 0.05], *P* = 0.026) and the choosiness dataset (*Zr* = -0.04 [-0.06, -0.01], *P* = 0.004) but not the mating success dataset (*Zr* = -0.001 [-0.02, 0.02], *P* = 0.86; Supplementary Figure 4). Similarly, the rank correlation test indicated significant variation in effect size over time for mating latency (*ρ* = 0.53, *P* = 0.003) and choosiness (*ρ* = -0.59, *P*<0.001), but no overall trend for mating success (*ρ* = 0.03, *P* = 0.81).

**Figure 3.**
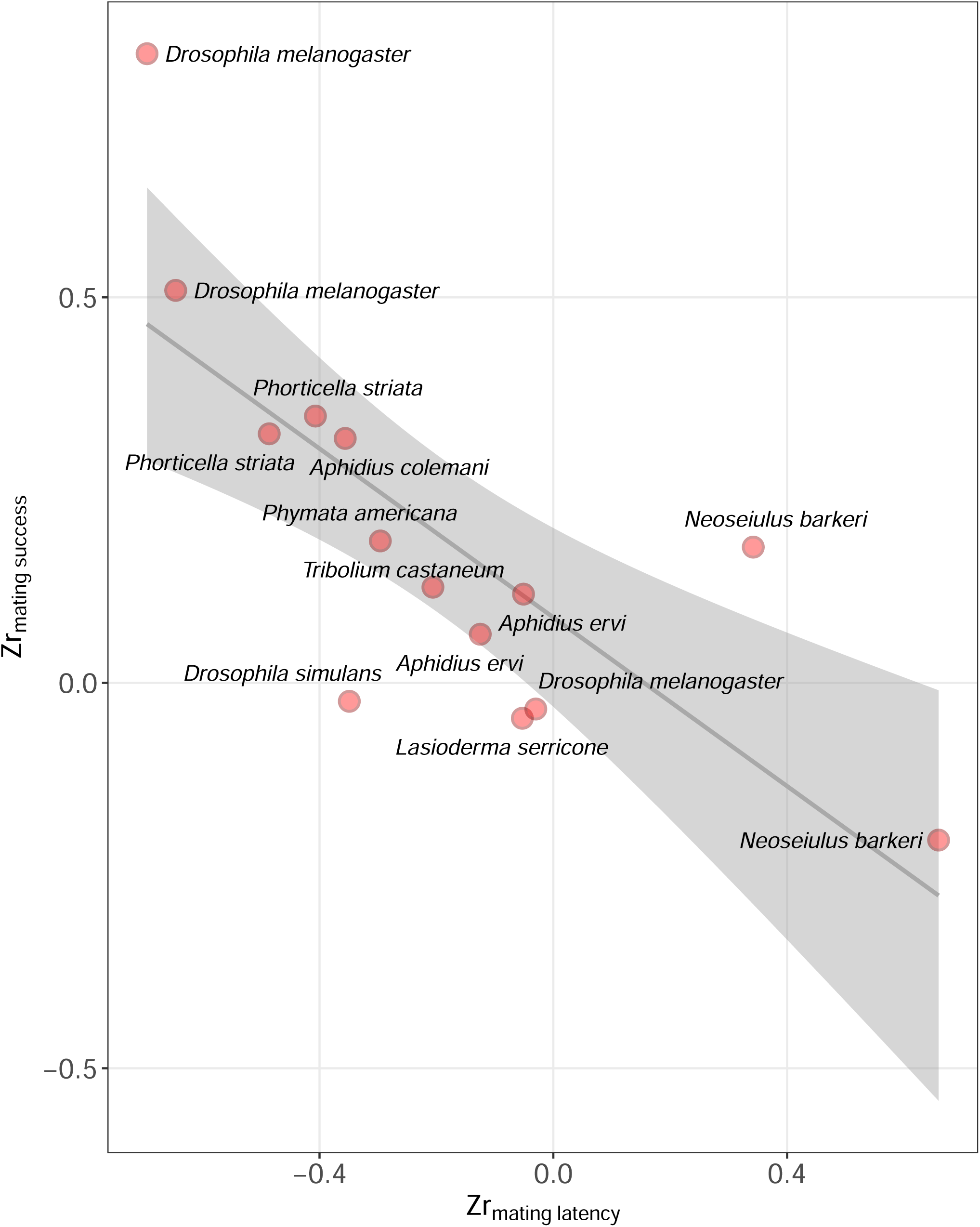
Fitted line and individual data points showing the relationship between *Zr*_mating latency_ and *Zr*_mating success_ for studies that measured both traits (14 effect sizes from 10 studies on 9 different species). The shaded area around the line of best fit indicates the 95% confidence interval.

### Main effects models

Phylogeny failed to resolve any heterogeneity in our main effects models; the estimates from phylogenetic and non-phylogenetic models were very similar or identical. Our intercept-only models for mating latency (non-phylogenetic method: *Zr* = -0.13 [-0.32, 0.05], phylogenetic method: *Zr* = -0.13 [-0.31, 0.05]), choosiness (non-phylogenetic method: *Zr* = 0.001 [-0.17, 0.18], phylogenetic method: *Zr* = 0.001 [-0.17, 0.18]), and mating success (non-phylogenetic method: *Zr* = 0.06 [-0.11, 0.22], phylogenetic method: *Zr* = 0.06 [-0.11, 0.22]) showed no overall effect of temperature (Figure 1; Supplementary Figures 5 & 6).

### Moderator models

As in the main effects models, the estimates from the phylogenetic and non-phylogenetic moderator models were similar or identical in all analyses. Statistical results reported below are based on the non-phylogenetic models (see Supplementary Table 2 for phylogenetic models).

There was some variation in the effect of temperature on mating latency depending on whether the animals were exposed to a temperature treatment during early development (*Zr* = 0.19 [-0.21, 0.59]), in adulthood before mating (*Zr* = -0.10 [-0.39, 0.19]), or during mating (*Zr* = -0.30 [-0.57, -0.03]), but this moderator (“time of temperature treatment”) was not statistically significant overall (QM=4.05, P=0.13, *R*^2^_marginal_=0.075; Figure 2). The effects of temperature on mating latency also did not vary in response to the type of temperature treatment (acute exposure: *Zr* = -0.17 [-0.40, -0.06]; acclimation: *Zr* = -0.06 [-0.41, 0.28]; QM=2.23, P=0.33) or the intensity of the treatment (*Zr* = -0.01 [-0.04, 0.01], QM=1.50, P=0.22).

The relationship between temperature and choosiness was not influenced by the time of the temperature treatment (early development: *Zr* = 0.05 [-0.15, 0.26]; during mating trial: *Zr* = -0.04 [-0.23, 0.14]; QM=1.32, P=0.25). Similarly, this relationship did not vary based on the type of temperature treatment (acute exposure: *Zr* = -0.05 [-0.26, 0.15]; acclimation: *Zr* = 0.02 [-0.17, 0.21; QM=0.84, P=0.66) or the intensity of the temperature treatment (*Zr* = -0.03 [-0.06, 0.01]; QM=2.59, P=0.11).

The effect of temperature on mating success varied depending on whether the animals were exposed to a temperature treatment during early development (*Zr* = -0.07 [-0.30, 0.17]), in adulthood before mating (*Zr* = -0.17 [-0.38, 0.04]), or during mating (*Zr* = 0.17 [0.01, 0.34]). Higher temperatures during the mating trial were associated with a higher mating success (QM=15.6, P<0.001; Figure 2). This moderator (“time of temperature treatment”) explained about 5% of the heterogeneity among effect sizes (*R*^2^_marginal_=0.048). In contrast, the effects of temperature on mating latency did not vary in response to the type of temperature treatment (acute exposure: *Zr* = 0.09 [-0.11, 0.29]; acclimation: *Zr* = -0.09 [-0.29, 0.11]; QM=0.89, P=0.64) or the intensity of the treatment (*Zr* = 0.01 [-0.02, 0.03], QM=0.36, P=0.55). Lastly, there were no substantial differences in effect sizes between studies that exposed males (*Zr* = -0.21 [-0.59, 0.17), females (*Zr* = -0.19 [-0.70, 0.33]), or both sexes (*Zr* = 0.09 [-0.10, 0.27]) to the temperature treatment (QM=3.14, P=0.37).

### Relationship between *Zr*_mating latency_ and *Zr*_mating success_

We found a strong negative correlation between the effect sizes (*Zr*) for mating latency and mating success from studies that measured both traits (*r*=-0.77, *t*_*12*_=-4.13, P=0.001; Figure 3). Similarly, our meta-regression showed that *Zr*_mating latency_ explained 75% of the heterogeneity in *Zr*_mating latency_ (QM=18.6, P<0.0001, *R*^2^_marginal_=0.745). This negative relationship indicates that in studies where a higher temperature led to an increase in mating latency, this was associated with a decrease in mating success, and vice versa (Figure 3).

## DISCUSSION

Our meta-analysis of 53 studies found no evidence for a global effect of temperature on mating latency, choosiness, or mating success. There was an increase in mating success when animals were exposed to higher temperatures during mating, but not when they were exposed before mating. Interestingly, in a subset of studies that measured both mating latency and mating success, we found a strong negative relationship between the effect sizes for these traits. This suggests that a decrease in mating latency at higher temperatures was associated with an increase in mating success.

These temperature-induced changes in mating behaviour may be mediated through physiological changes (Abram et al. 2017). The body temperature and metabolic rate of ectothermic animals is directly influenced by changes in ambient temperature, which can affect their activity levels (Kearney et al. 2010, Gunderson & Leal 2015). This also means that higher temperatures might lead to increased costs of mate searching and mate assessment (Punzalan et al. 2008, García-Roa et al. 2020). For example, ambush bugs (*Phymata americana*) have reduced mate-searching success at higher ambient temperatures (Punzalan et al. 2008), which might result in a higher propensity to mate with a potential partner under these conditions.

Previous work has suggested that temperature may also affect mating behaviour through changes in communication between mates (Martin & Lopez 2013). Both mating latency and mating success are dependent of the ability of a male to stimulate the female for mating. When mate communication involves chemical signals over large areas, such as femoral secretions to mark territories, females may take longer to detect them due to faster evaporation at higher temperatures (Martin & Lopez 2013). Such a disruption in mate communication may lead to a higher mating latency and lower mating success in warmer environments. On the other hand, when mate communication involves chemical signals at a small spatial scale, a higher temperature may increase the volatility of these chemicals and improve mate recognition and assessment. This may lead to a shorter mating latency and higher mating success in warmer environments. In line with this latter scenario, our meta-analysis found evidence for an increase in mating success when animals were exposed to higher temperatures during mating, but it is unclear whether the underlying mechanism for this pattern was a change in mate communication through chemical signals.

Contrary to our expectation, we found no evidence that temperature-induced changes in mating success were accompanied with changes in mating latency or choosiness. This was somewhat surprising, given that mating latency, choosiness, and mating success are closely linked (Lindström & Lehtonen 2013, Breedveld & Fitze 2015). For example, a choosier individual may require more time for mate assessment, leading to an increase in mating latency, which can then result in lower mating success (Hegde & Krishna 1997). Similarly, a choosier individual may be less likely to mate with low-quality partners, resulting in lower mating success. The absence of a directional effect of temperature on mating latency and choosiness, despite its effects on mating success, may be partly due to the use of different datasets for each of these traits (Table 1). In fact, only one of the studies in our meta-analysis included data on all three traits (Supplementary Table 2). In a subset of studies that measured both mating latency and mating success, there was a strong negative relationship between the effect sizes for these traits, as we had expected (Figure 3). This supports our interpretation that the different patterns we observed in mating latency, choosiness, and mating success may be due to differences in the species used across the three datasets. A recent review argued that the relationship between temperature and sexual selection is likely to vary across species in relation to their mating system, physiology, and behaviour (García-Roa et al. 2020). The absence of global directional effects in mating behaviour in our meta-analysis provides support for this prediction. Importantly, our findings suggest that due to substantial among-species variation, it may be difficult to generate predictions for how the strength of sexual selection in natural populations will change in a warming world.

Phylogeny did not seem to influence the effects of temperature on mating behaviour across the range of species included in our analysis (mating latency: *n* = 14, choosiness: *n* = 14, choosiness: *n* = 28). This may be because certain features of our datasets make the detection of a phylogenetic signal unlikely. For example, mating behaviour has the capacity to evolve rapidly and is thus evolutionarily labile (Blomberg et al. 2003, Dougherty & Shuker 2015). Our analysis also includes measures of mating preference (choosiness) for a wide range of traits, such as body size, colouration, and mating calls, which might make it more difficult to find a phylogenetic signal. Lastly, it is worth noting that a large majority of studies included in our analysis were on insects (Table 1), limiting our ability to draw conclusions about general patterns across taxa. For example, we initially intended for our meta-analysis to include a comparison between endotherms and ectotherms. We expected that temperature changes would have a stronger effect on mating behaviour and mating success in ectotherms, given that ambient temperature can directly affect their body temperature, metabolic rate, locomotor performance, and activity levels (Gibert et al. 2007, Kearney et al. 2010, Lachenicht et al. 2010, Gunderson & Leal 2015). However, it was not possible to carry out this comparison due to the relevant studies available in published literature. Out of 53 studies included in our meta-analysis, only two were on endotherms (birds). We therefore strongly encourage future research on the effects of temperature on mating behaviour and mating success in endotherms, as well as ectotherms other than insects.

Our publication bias tests suggest there may be some influence on the overall results. Firstly, there was evidence for funnel asymmetry for mating success, suggesting a potential for publication bias in this dataset. The observed bias may have also been caused by unexplained heterogeneity among studies due to other moderators that we did not consider in our analysis. Secondly, we found evidence for a time-lag bias in the mating latency and choosiness datasets, where there was a trend for a decrease in effect size over time (Supplementary Figure 4). This is a common pattern in meta-analyses in ecology and evolutionary biology (Jennions and Møller 2002). We have not used methods such as trim and fill that try to compensate for publication bias, as they may perform poorly in high heterogeneity datasets (Moreno et al., 2009, Moran et al. 2020). Regardless of the underlying causes for this publication bias, it is important to take it into consideration when interpreting the results, particularly in cases where confidence intervals are close to zero, such as the effect of temperature on mating success (Figure 2b).

Another limitation of our study was that the sample size for the mating latency (*k*=29) and choosiness datasets (*k*=29) was relatively small. As a result, subset analyses testing for the effects of moderators had small sample sizes for each factor (Supplementary Table 1). Most of these analyses did not detect any significant effects, with the exception of an effect of the time of temperature treatment on mating success, which was discussed above. The fact that the intensity and duration of the temperature treatment did not have an overall effect on mating behaviour and mating success was surprising and may be due to low statistical power.

Our meta-analysis focused on the effects of temperature on mating success and two precopulatory traits, mating latency and choosiness. Nevertheless, temperature can also influence postcopulatory processes (García-Roa et al. 2020). For example, the amount and quality of sperm transferred during mating has been shown to vary with temperature (Reinhardt et al. 2015, Sales et al. 2018, Gasparini et al. 2018, Walsh et al. 2019). A study on the cigarette beetle (*Lasioderma serricone*) that examined both precopulatory and postcopulatory traits actually found that temperature had a stronger effect on the latter (Suzaki et al. 2018). We therefore suggest that a meta-analysis on how temperature variation also influences postcopulatory traits, such as sperm production and sperm competition, may be worthwhile.

Given rising temperatures due to global climate change, it is important to better understand how changes in temperature may affect mating patterns and sexual selection (García-Roa et al. 2020). Here, we show an increase in mating success when animals were exposed to higher temperatures during mating, but not during early development or in adulthood before mating. We found no evidence for directional effects of temperature on mating latency or choosiness, suggesting it may be difficult to generate general predictions for how the strength of sexual selection will change in a warming world. Nevertheless, we also found a strong negative relationship between the effect sizes for mating latency and mating success. This suggests that in species where a higher temperature leads to an increase in mating latency, this may result in a decrease in mating success, and vice versa. Our meta-analysis therefore provides new insights into the effects of temperature on mating behaviour and sexual selection.

## ACKNOWLEDGMENTS

We thank the Associate Editor and three anonymous reviewers for their helpful comments on an earlier version of this manuscript.

## AUTHOR CONTRIBUTIONS

AB carried out the data extraction, contributed to the data analysis, and provided comments on the manuscript draft. NP conceived the study, contributed to data extraction and data analysis, and wrote the manuscript.

## DATA AVAILABILITY STATEMENT

All data used in this meta-analysis and the associated R code will be archived on the Dryad Digital Repository upon manuscript acceptance.

