## Supplementary for "Effects of temperature on mating behaviour and mating success: a meta-analysis"

**Supplementary Table 1.** Number of effect sizes (*k*) in our data sets on mating latency, choosiness, and mating success.

| | Mating latency | Choosiness | Mating success |
|---|---|---|---|
| *Sex* | | | |
| Female | 19 | 22 | 42 |
| Male | 0 | 5 | 2 |
| Both | 10 | 2 | 14 |
| *Time of temperature treatment* | | | |
| Early development | 6 | 13 | 6 |
| Before mating | 11 | 0 | 14 |
| During mating | 12 | 16 | 38 |
| *Type of temperature treatment* | | | |
| Acute exposure | 20 | 9 | 29 |
| Acclimation | 9 | 20 | 29 |

**Supplementary Table 2.** Summary of data extracted from each study used in the meta-analysis. For each study, we present an abbreviated reference to the study, the scientific name of the study species, and the types of data extracted from that study (i.e., mating latency, choosiness, and/or mating success).

| **Study** | **Species** | **Mating latency** | **Choosiness** | **Mating success** |
|---|---|---|---|---|
| Albrecht *et al.* (1999) | *Pomacea canaliculate* | x | | |
| Amin *et al.* (2010) | *Bombus terrestris* | | | x |
| Arbogast (2007) | *Plodia interpunctella* | | | x |
| Beaulieu & Sockman (2012) | *Melospiza lincolnii* | | x | |
| Beckers & Schul (2008) | *Neoconocephalus triops* | | x | |
| Brandt *et al.* (2018) | *Habronattus clypeatus* | | | x |
| Caetano & Hajek (2017) | *Sirex noctilio* | | | x |
| Colinet & Hance (2009) | *Aphidius colemani* | x | | x |
| Conrad *et al.* (2017) | *Osmia bicornis* | | | x |
| Coomes *et al.* (2019) | *Taeniopygia guttata* | | x | |
| Delisle (1995) | *Choristoneura rosaceana* | | | x |
| Dubey *et al.* (2016) | *Menochilus sexmaculatus* | | x | |
| Everman *et al.* (2018) | *Drosophila melanogaster* | x | | x |
| Fasolo & Krebs (2004) | *Drosophila melanogaster, Drosophila simulans, Drosophila mojavensis* | | | x |
| Geister & Fischer (2007) | *Bicyclus anynana* | | | x |
| Gerhardt (2005) | *Hyla chrysoscelis* | | x | |
| Goebel (2006) | *Chilo sacchariphagus* | | | x |
| Grace & Shaw (2004) | *Laupala cerasina* | | x | |
| Hsu & Wu (2001) | *Ctenocephalides felis* | | | x |
| Ingleby *et al.* (2013) | *Drosophila simulans* | x | | x |
| Ismail *et al.* (2010) | *Aphidius ervi* | x | x | x |
| Janowitz & Fischer (2011) | *Bicyclus anynana* | x | | |
| Jiao *et al.* (2009) | *Pardosa astrigera* | x | | |
| Kindle *et al.* (2006) | *Gryllodes sigillatus, Acheta domestricus* | | | x |
| Kvanermo & Forsgren (2000) | *Pomatoschistus minutus* | | x | |
| Laudien & Seifert (1983) | *Drosophila simulans* | | | x |
| McKibben & Bass (1998) | *Porichthys notatus* | | x | |

| | | | | |
|---|---|---|---|---|
| Mhatre *et al.* (2011) | *Oecanthus henryi* | x | | |
| Milner *et al.* (2010) | *Uca mjoebergi* | | x | |
| Olvido *et al.* (2010) | *Allonemobius socius* | | x | |
| Papadopoulou (2006) | *Lasioderma serricone* | | | x |
| Parkash *et al.* (2011) | *Drosophila melanogaster* | x | | |
| Patton & Krebs (2001) | *Drosophila melanogaster, Drosophila simulans, Drosophila mojavensis* | | | x |
| Pires & Hoy (1992) | *Gryllus firmus* | x | | |
| Punzalan *et al.* (2008) | *Phymata americana* | x | | x |
| Putz & Crews (2005) | *Eublepharis maculatus* | | x | |
| Ritchie *et al.* (2001) | *Drosophila montana* | | x | |
| Sambucetti & Norry (2015) | *Drosophila buzzatii* | | | x |
| Scharf *et al.* (2019) | *Tribolium castaneum* | x | | x |
| Sih *et al.* (2002) | *Aquarius remiges* | | | x |
| Singh *et al.* (2016) | *Drosophila melanogaster* | x | | x |
| Stazione *et al.* (2019) | *Drosophila melanogaster* | x | | |
| Suzaki *et al.* (2018) | *Lasioderma serricone* | x | | x |
| Symes *et al.* (2017) | *Oecanthus forbesi* | | x | |
| Vasudeva *et al.* (2018) | *Callosobruchus maculatus* | | | x |
| Westerman & Monteiro (2016) | *Bicyclus anynana* | x | | |
| Wilson *et al.* (2007) | *Gambusia holbrooki* | | | x |
| Yang *et al.* (2017) | *Nilaparvata lugens* | | | x |
| Yenisetti *et al.* (2006) | *Phorticella striata* | x | | x |
| Zhang *et al.* (2013) | *Plutella xylostella* | x | | x |
| Zhang *et al.* (2016) | *Neoseiulus barkeri* | x | | |
| Zizzari & Ellers (2011) | *Orchesella cincta* | | | x |
| Zverev *et al.* (2018) | *Chrysomela lapponica* | | | x |

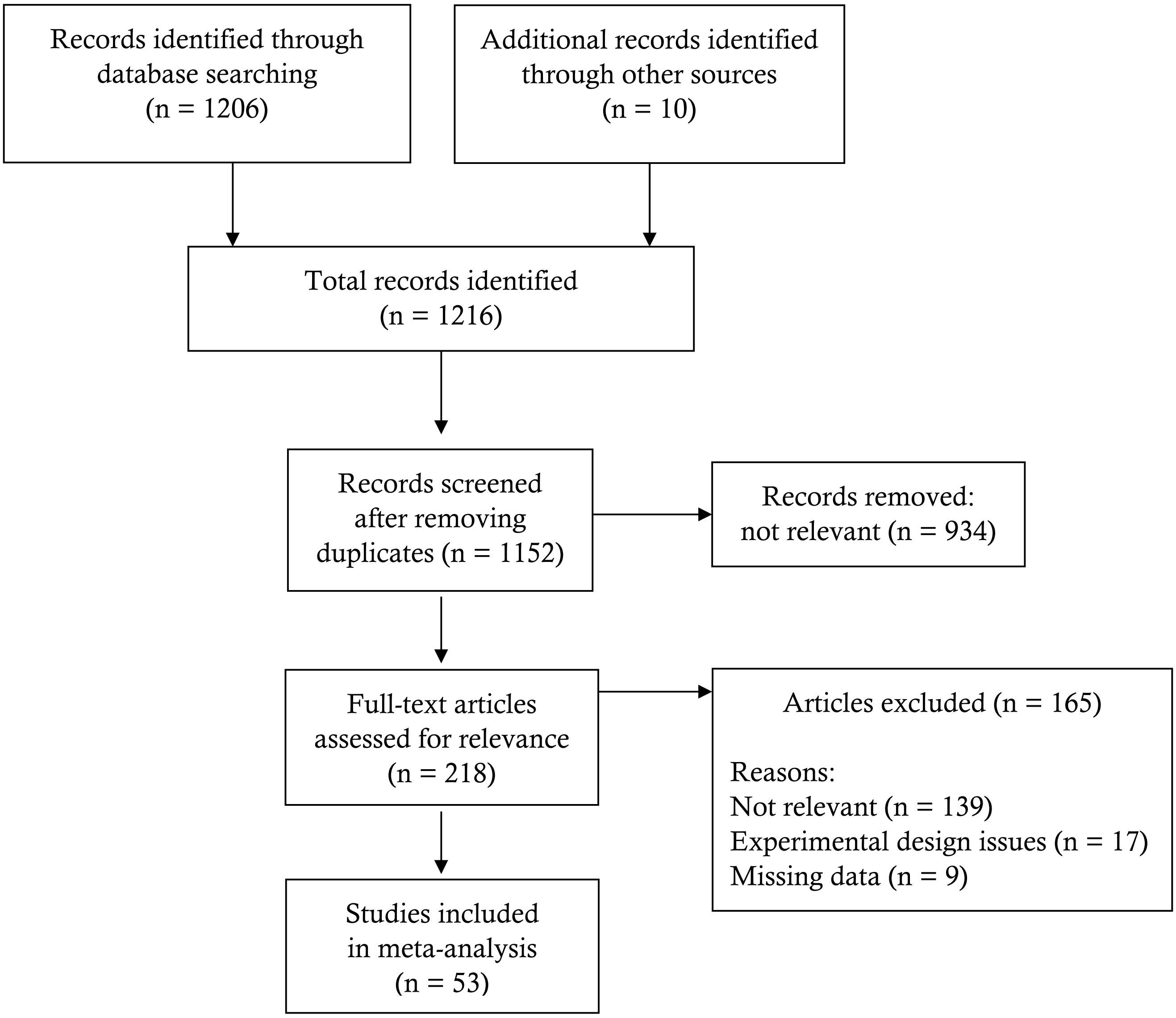


**Supplementary Figure 1.** PRISMA diagram showing the selection process for the studies included in this meta-analysis.

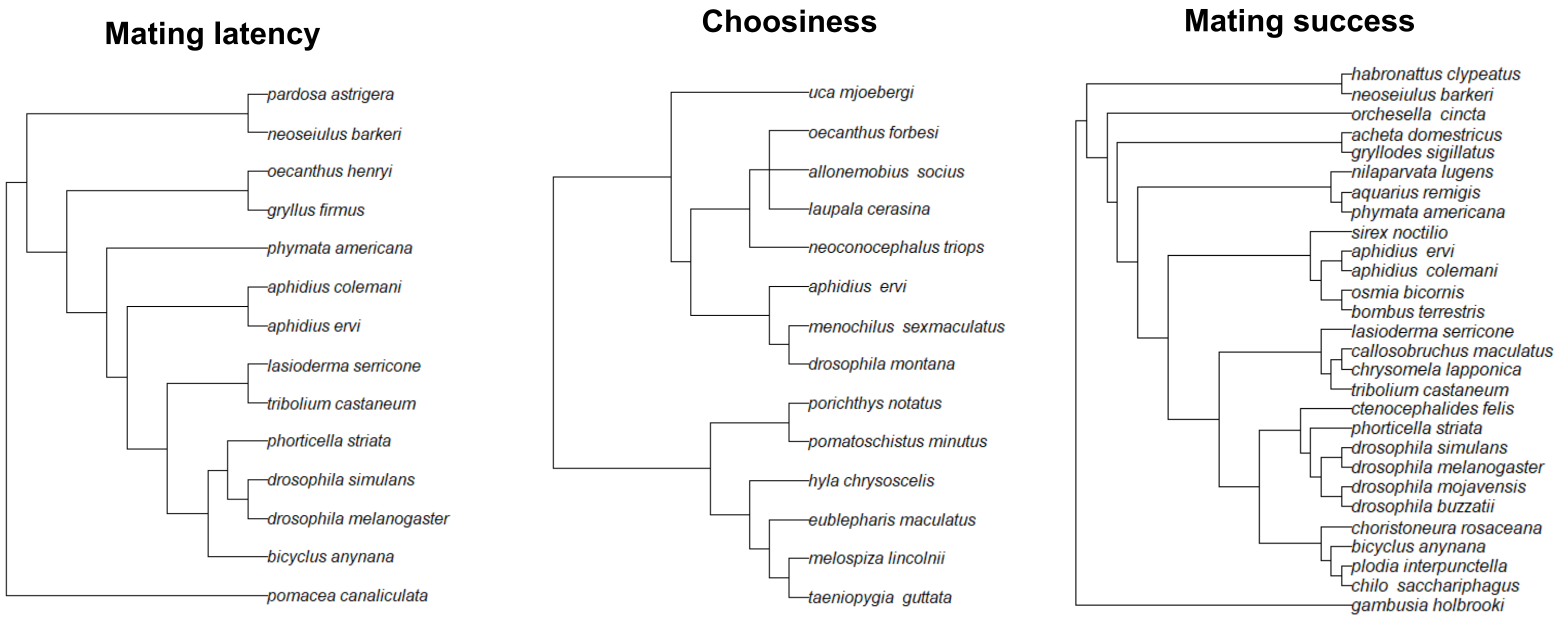


**Supplementary Figure 2.** Phylogenetic trees for the species included in the mating latency, choosiness, and mating success datasets used in the meta-analysis.

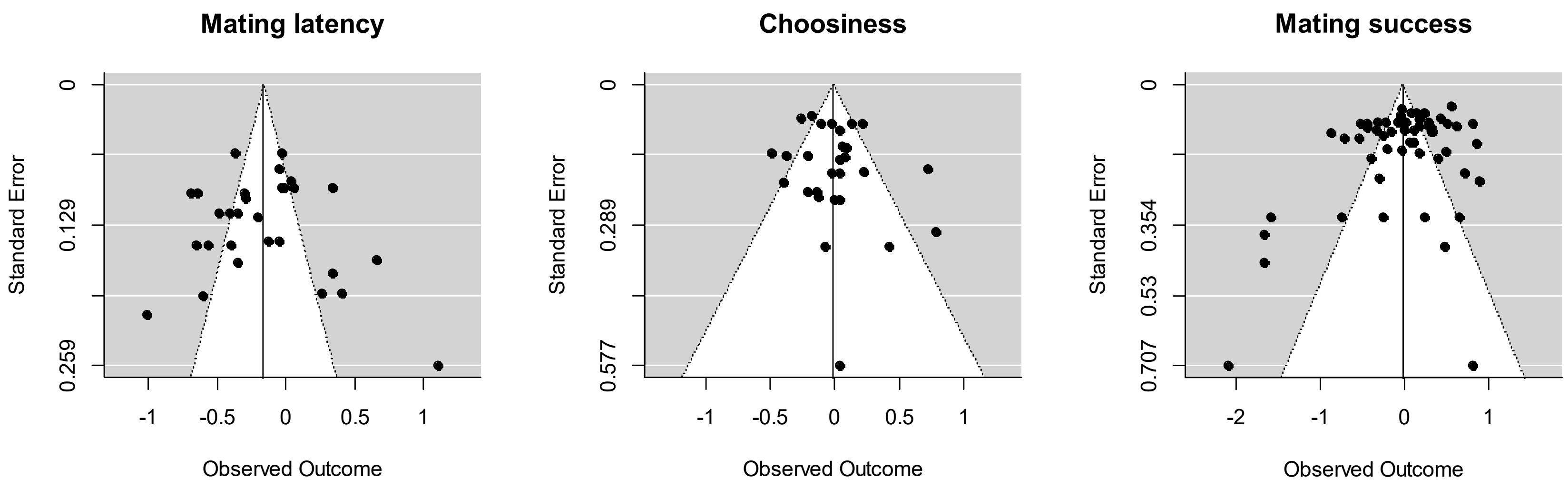


**Supplementary Figure 3.** Funnel plots generated to examine the potential for underreporting of non-significant results in each of our three datasets (mating latency, choosiness, and mating success).

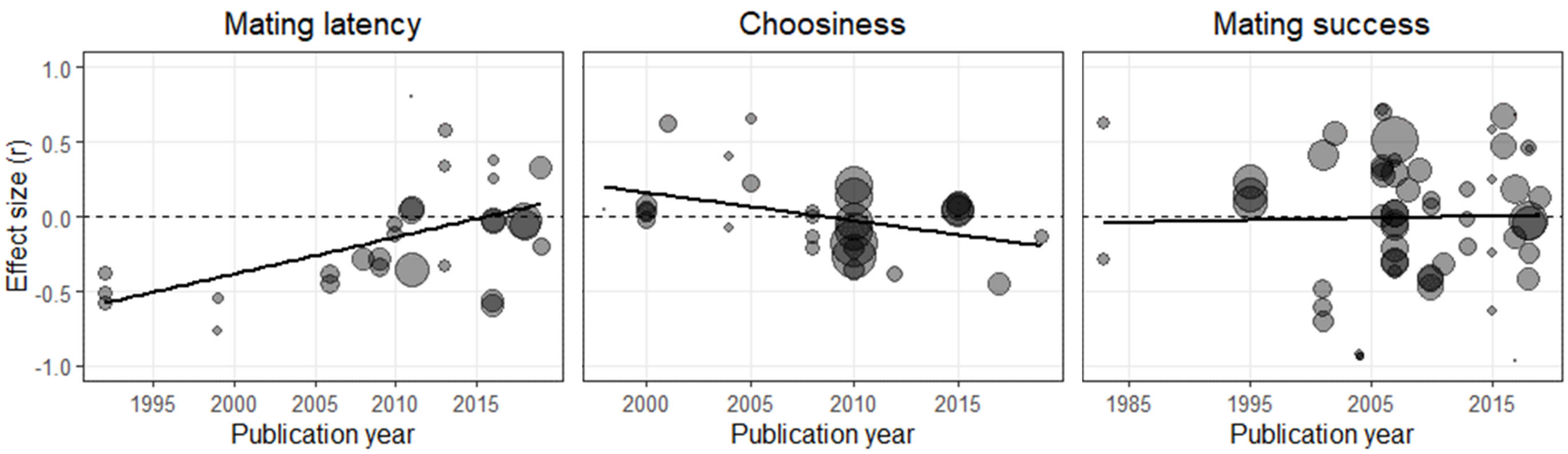


**Supplementary Figure 4.** Effect size (r) of the relationship between temperature and mating latency, choosiness, or mating success over time. The relative size of each point represents the sample size of each effect size. The solid black line represents the regression of effect size by publication year.

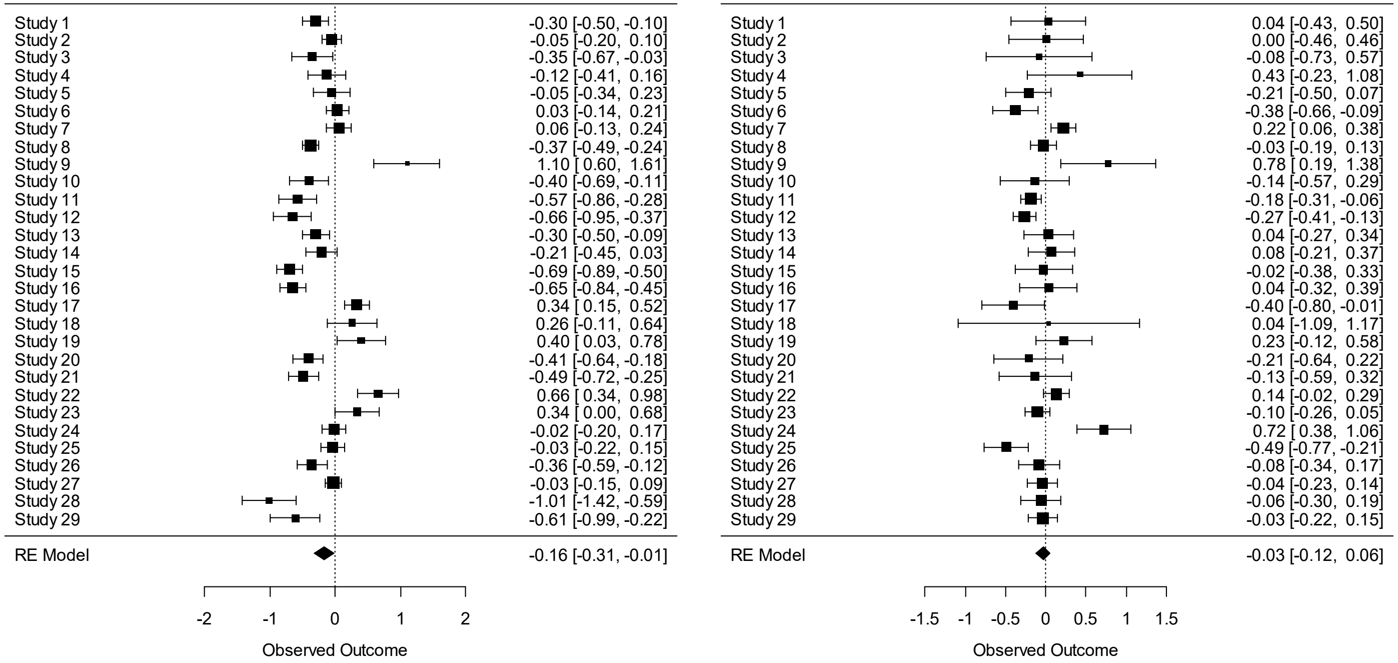


**Supplementary Figure 5.** Forest plots for mating latency (left) and choosiness (right).

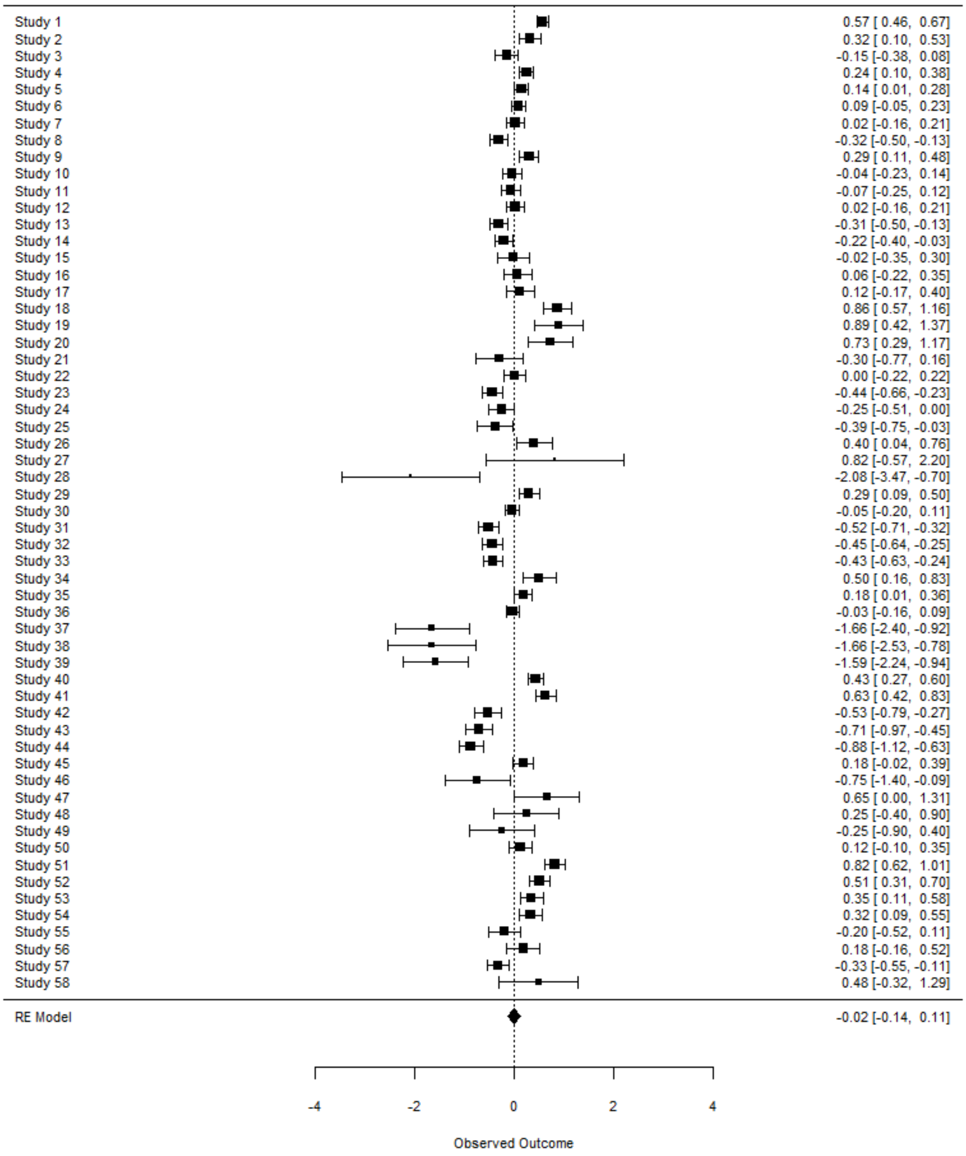


**Supplementary Figure 6.** Forest plot for mating success.
